# An enhanced inverted encoding model for neural reconstructions

**DOI:** 10.1101/2021.05.22.445245

**Authors:** Paul S. Scotti, Jiageng Chen, Julie D. Golomb

## Abstract

Here we present a more interpretable and versatile approach for reconstructing the contents of perception, attention, and memory from neuroimaging data. Our enhanced inverted encoding model (eIEM) incorporates theoretical and methodological improvements including proper accounting of population-level tuning functions and a trial-by-trial prediction error-based metric where reconstruction quality is measured in meaningful units. Added functionality and improved flexibility is further gained via eIEM’s novel goodness-of-fit feature: for trial-by-trial reconstructions, goodness-of-fits are obtained independently (non-circularly) to prediction error and can be applied to any IEM procedure or decoding metric, resulting in improved reconstruction quality and brain-behavior correlations, and more creative applications. We validate eIEM from methodological principles, simulated neuroimaging datasets, and three pre-existing fMRI datasets spanning perception, attention, and working memory. Notably, eIEM is easy to apply and broadly accessible – our Python package (https://pypi.org/project/inverted-encoding) implements eIEM in one line of code – and is easily modifiable to compare performance metrics and/or scale up to more complex models.

## 1. Introduction

A mental representation can be defined as the “systematic relationship between features of the natural world and the activity of neurons in the brain”^1^. An increasingly common approach to study mental representations using neuroimaging data is to employ encoding models, which describe this relationship computationally, typically by reducing the complexity of the input data with a set of functions that, when combined, roughly approximate the neural signal.

Encoding and decoding models (aka voxelwise modeling or stimulus-model based modeling) have become a standard method for investigating neural representational spaces and predicting stimulus-specific information from brain activity^2–4.^. The key advantages of such models over other computational approaches such as multivariate pattern classification or representational similarity analysis are typically touted as the following: (1) Encoding models can take inspiration from single-unit physiology by consisting of tuning functions in stimulus space (aka feature space), allowing both the maximally receptive feature and the precision/sensitivity of the tuning to be estimated across a population of neurons. (2) Encoding models (which transform presented stimuli into predicted brain activity) are easily inverted into decoding models (which transform observed brain activity into predicted stimuli), and this process can be applied to a wide range of stimuli, including oriented Gabor patches^5, 6^, colors^7^, acoustic musical features^8^, and human faces^9^. (3) The decoding model can predict, or *reconstruct*, novel stimuli or experimental conditions not used in the training of the model, with reconstructions offering interpretational advantages beyond classification-based decoding by yielding measures of activity across the full stimulus space and facilitating research questions exploring representational schemes for brain regions of interest^4, 7^.

The inverted encoding model (IEM), sometimes also referred to as forward modeling, is one example of an encoding and decoding model that has quickly risen to prominence in the cognitive neuroscience community (see Supplemental Figure 1)^10–36^. The basic idea behind IEMs is illustrated in Figure 1A. First, an encoding model is trained to associate patterns of brain activity with specific stimuli. The model uses simple linear regression and a basis set representing the hypothesized population-level tuning functions, consisting of several channels that are modeled as cosines (or von Mises functions) equally separated across stimulus space (e.g., orientation, color, spatial location). Each channel in the basis set can be assigned a weight per voxel (we adopt fMRI nomenclature here, but IEMs can be applied to any modality like EEG, MEG, etc.) and hence a model can be trained to predict the activity of each voxel using the weights of each channel as predictors (i.e., the regressors in a linear regression). Then, this trained encoder is inverted such that it becomes a decoder capable of reconstructing a trial’s stimulus when provided with a novel set of voxel activations (see Methods for mathematical implementation).

**Figure 1.**
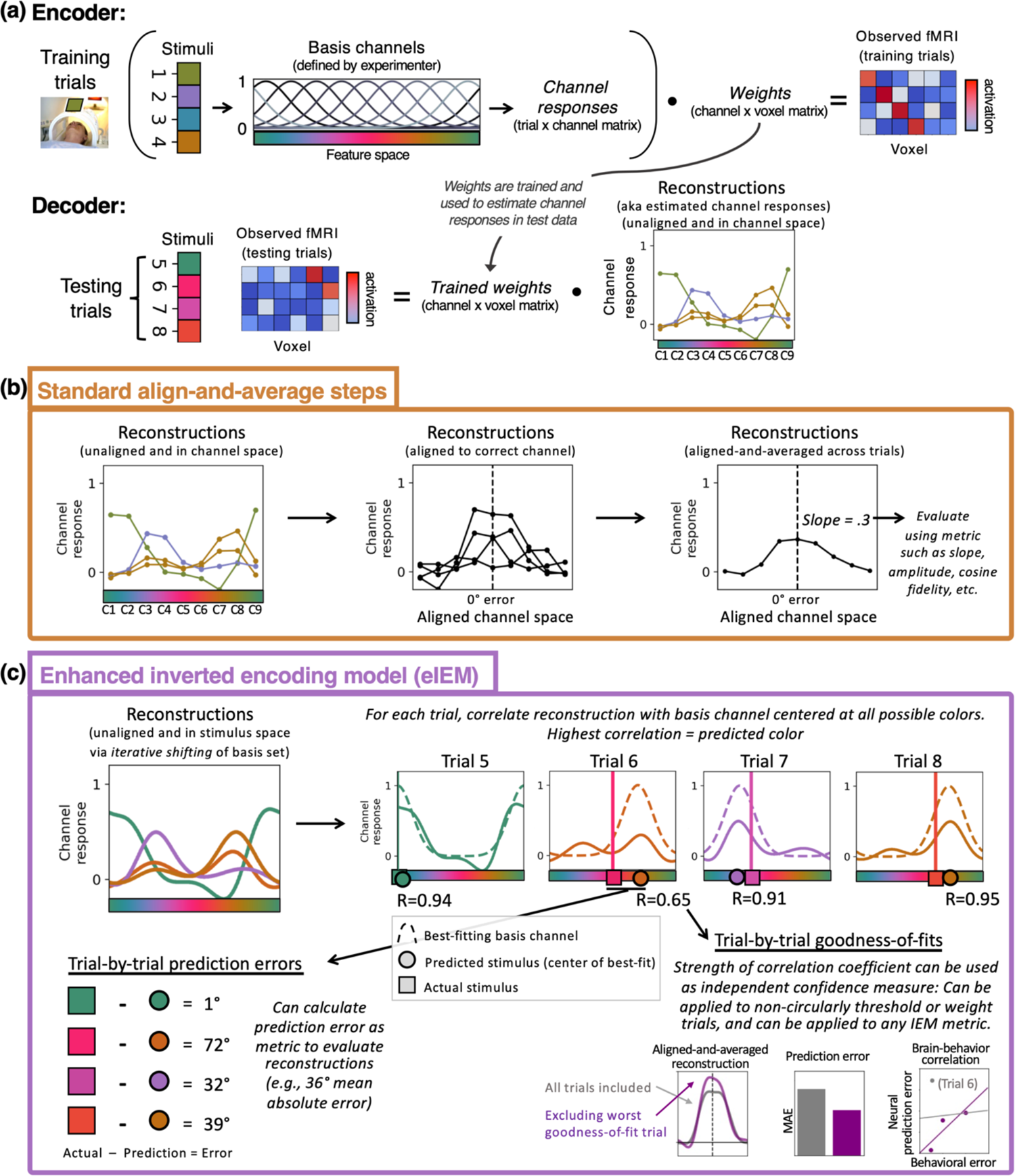
An overview of the steps involved in eIEM using a toy example of an fMRI experiment reconstructing stimulus colors. (a) Basic encoder and decoder steps common to all variations of IEM. The prerequisites for implementing an IEM are the feature value of the stimulus for every trial (here 8 trials of colored squares), a trial-by-voxel matrix of brain activations for every voxel per trial (here simulated beta weights from a six-voxel brain region, and a basis set representing hypothesized population-level tuning functions; see Methods for details). The encoder models each voxel’s response as the weighted sum of the channels. Given that the trial-by-voxel matrix and the basis set are already given, the weights matrix can be estimated via least-squares linear regression. Once the weights matrix is estimated from the training dataset, it can be inverted such that the encoder becomes a decoder for the test dataset. Now, instead of estimating the weights matrix via least-squares linear regression, the weights matrix and the trial-by-voxel matrix are given and the channel responses (i.e., reconstructions) are estimated. The resulting estimated channel responses, or simply *reconstructions*, is a trial-by-channel matrix where each trial has its own reconstruction composed of weighted cosines. For simplicity, this example shows the encoder trained on the first half of trials and the decoder used to predict the color of the remaining trials, but in most applications cross-validation should be used such that every trial may be decoded while avoiding circularity/double-dipping. (b) The commonly applied align-and-average procedure, which involves aligning and averaging reconstructions across trials and measuring the result according to a variety of possible metrics (e.g., amplitude, slope; see Supplemental Figure 1). (c) eIEM deviates from this step by evaluating trial-by-trial reconstructions rather than an averaged reconstruction. We use iterative shifting of the basis set to allow channel space to equal stimulus space, correlate the reconstructions with the full set of basis channels to estimate each trial’s predicted stimulus (and then average prediction error), and also estimate goodness-of-fits for each trial’s reconstruction.

IEM is popular due to its simplicity, robust performance, and grounding in single-unit physiology principles. Over recent years, several variations, improvements, and additional features have been proposed for IEMs, resulting in various approaches and evaluation metrics (Supplemental Figure 1). This malleability is both an advantage and a challenge of IEM, as there is sometimes ambiguity over which metric is best, particularly for less experienced users. Moreover, as discussed more below, certain implementations of IEM can produce misleading or difficult to interpret results, and do not account for the shape of the basis channels (as recently pointed out in some high-profile debates in the literature^37–40)^. While this is not to suggest that previous papers have made incorrect scientific interpretations, there are several aspects of IEM that can be improved upon.

Here we propose a variation of IEM which improves the interpretability of stimulus reconstructions, addresses some key issues inherent in the IEM procedure as commonly applied, and provides trial-by-trial stimulus predictions and goodness-of-fit estimates. Our proposed approach serves two main functions: first, it can be thought of as a set of recommended best-practices based on theoretical principles, and second, it offers some novel enhancements to the IEM procedure, including a focus on trial-by-trial predictions that includes a built-in measure of uncertainty. For simplicity and ease of communication, we refer to this approach as “enhanced inverted encoding modeling” (eIEM). In this manuscript we provide theoretical justification for each of the recommended aspects of eIEM, along with real and simulated data confirming that eIEM produces results at least as good as – and in several cases better than – a commonly implemented alternative. We also share a publicly available Python package that can apply eIEM to neuroimaging data in a single line of code. While advanced researchers may have reason for adopting some aspects of eIEM without others (using either our code or their own), more novice users may find this pre-packaged set of recommendations and enhanced functionality particularly useful.

Figure 1 depicts the eIEM procedure. In essence, eIEM is a combination of several modifications and improvements (some previously employed, some novel). First, with eIEM, the core encoder and decoder steps described above (i.e., least-squares linear regression for estimating channel weights and responses, using a basis set of hypothesized population-level tuning functions) remain the same, but the encoding model is repeatedly fitted with slightly shifted basis sets such that subsequent reconstructions are in stimulus space rather than channel space. This “iterative shifting” of the basis set has been employed in a few previous papers^41–43^, but it is not yet common practice. Iterative shifting allows for more accurate reconstructions spanning the full stimulus space (allowing the remaining eIEM steps to be optimally applied), and aids more generally in reducing potential interpretation flaws and bias associated with impoverished basis sets, as illustrated and explained in Supplemental Figure 2. We also include a Jupyter notebook that illustrates the advantages of iterative shifting across various simulated neuroimaging data on OSF (https://osf.io/vf3n6).

More critically, the core difference of eIEM is that we analyze reconstructions at the trial-by-trial level, using a procedure that can obtain stimulus predictions, prediction error, and goodness-of-fits for each trial. In contrast, most existing implementations of IEM typically employ an align-and-average approach (Figure 1B), where each trial’s reconstruction is circularly shifted along the x-axis such that it is centered on the “correct” channel (closest to the ground truth stimulus feature), and then these aligned reconstructions are averaged together to result in a single reconstruction, typically quantified using metrics such as slope, amplitude, etc. As we show in the Results, this align-and-average approach is susceptible to information loss, outlier bias, and interpretation issues.

With eIEM, these aligned-and-averaged reconstructions can still be obtained for visualization purposes, but eIEM offers several novel advantages for evaluating, quantifying, and interpreting reconstructions. eIEM obtains trial-by-trial stimulus predictions using a correlation table approach^a^ (previously used in a small number of papers^7, 42^) that adapts to the shape of the basis channel (Figure 1C). For each trial, a set of correlation coefficients is computed, each reflecting the correlation between that trial’s reconstruction and a basis channel (i.e., “perfect reconstruction”) centered at every integer in stimulus space (e.g., resulting in 360 correlation coefficients for a stimulus space ranging from 0-359°). The highest of these correlation coefficients is determined to be the best fit for that trial, and the predicted stimulus feature for that trial is simply the center of that best-fitting basis channel. As described below in the Results, evaluating reconstructions using stimulus predictions obtained via this approach is more methodologically sound than, say, taking the location of highest amplitude as the stimulus prediction^25, 44^, because the shape of the basis functions is automatically taken into account.

To quantify reconstruction quality, eIEM naturally lends itself to prediction error based summary metrics. The difference between the predicted and actual stimulus feature is that trial’s prediction error; this error can be averaged across trials to obtain mean absolute error (MAE), mean signed error, etc. Here we adopted MAE (specifically circular mean absolute error when using circular feature spaces) as a default metric for evaluating reconstruction quality, because of its simplicity and grounding in meaningful units (average prediction error), particularly compared to other IEM metrics such as slope, amplitude, reconstruction fidelity, etc which use arbitrary units. However, MAE itself is not a necessary component of eIEM — trial-by-trial prediction errors can be evaluated in any manner the researcher sees fit.

The final and most novel advantage that eIEM offers is the automatic calculation of trial-by-trial goodness-of-fit values. Recall that each trial’s stimulus prediction is determined as the center of the best-fitting basis function. Because this is calculated independently of the actual correct stimulus, the goodness-of-fit values themselves (correlation coefficients) can also be recorded and optionally leveraged to estimate trial-by-trial confidence of predictions. Individual trials in a neuroimaging study can vary substantially in signal quality (driven by attentional fluctuations, alertness, head motion, scanner noise, etc.) but typically the IEM procedure does not incorporate uncertainty into decoding performance. The lack of uncertainty information has been noted in other contexts, with some recent alternatives to IEM proposed to incorporate uncertainty^45, 46^. eIEM has the advantage of easily and automatically producing a trial-by-trial measure of prediction uncertainty within the IEM framework itself. This enhancement adds substantial flexibility to the IEM procedure. For example, as depicted in Figure 1C, goodness-of-fit can be used to set thresholds in a non-circular manner, such that less reliable trials are excluded from reconstructions. As we demonstrate in the Results using real and simulated fMRI datasets, goodness-of-fit thresholding can substantially increase statistical power and performance in versatile ways. A caveat to this increased flexibility is that a researcher cannot try varying cutoff thresholds and then cherry-pick the threshold that provides desirable results; thresholds need to be determined before data collection or according to some sort of a priori, independent criteria.

Here we define eIEM as an encoding model that uses the combination of iterative shifting (or any technique which allows reconstructions to reside in stimulus space), evaluating reconstructions using the basis channel defined for the encoder (via the correlation table approach or other constrained model fitting), calculating trial-by-trial prediction errors, and calculating goodness-of-fits independently from stimulus predictions. Researchers can easily implement eIEM on their own through our publicly available Python package (https://pypi.org/project/inverted-encoding; see Methods for more information). Of course, researchers can choose to implement eIEM without our Python package or choose to only implement some but not all aspects of eIEM. The value of eIEM is primarily in terms of evaluating IEM results: using both real and simulated data, we show below that eIEM can be beneficial in terms of interpretability, flexibility, functionality, and robustness to methodological concerns.

## 2. Results

We first validate eIEM by implementing it on three existing fMRI datasets^43, 47, 48^ spanning the research topics of perception, attention, and memory. We then demonstrate the ways eIEM can go beyond the more standard IEM approaches, offering practical advantages and additional functionality. Finally, we use simulated reconstructions to illustrate some additional theoretical issues and methodological limitations that eIEM addresses. See the Methods for information regarding each dataset and how data were processed.

### 2.1 Validating eIEM on real fMRI data

First, as initial validation, we confirmed that across all three datasets eIEM replicated the overall pattern of results obtained previously with IEM (Figure 2). To do this in a rigorous and consistent way, we reanalyzed all datasets ourselves using a “standard” align-and-average IEM approach with slope as the decoding metric, even if the original paper did not employ this exact same procedure. We chose slope as it is a commonly used decoding metric across IEM papers (see Supplemental Figure 1). For the purposes of testing eIEM, we selected a single primary analysis from each of the three datasets, as detailed in the Methods.

**Figure 2.**
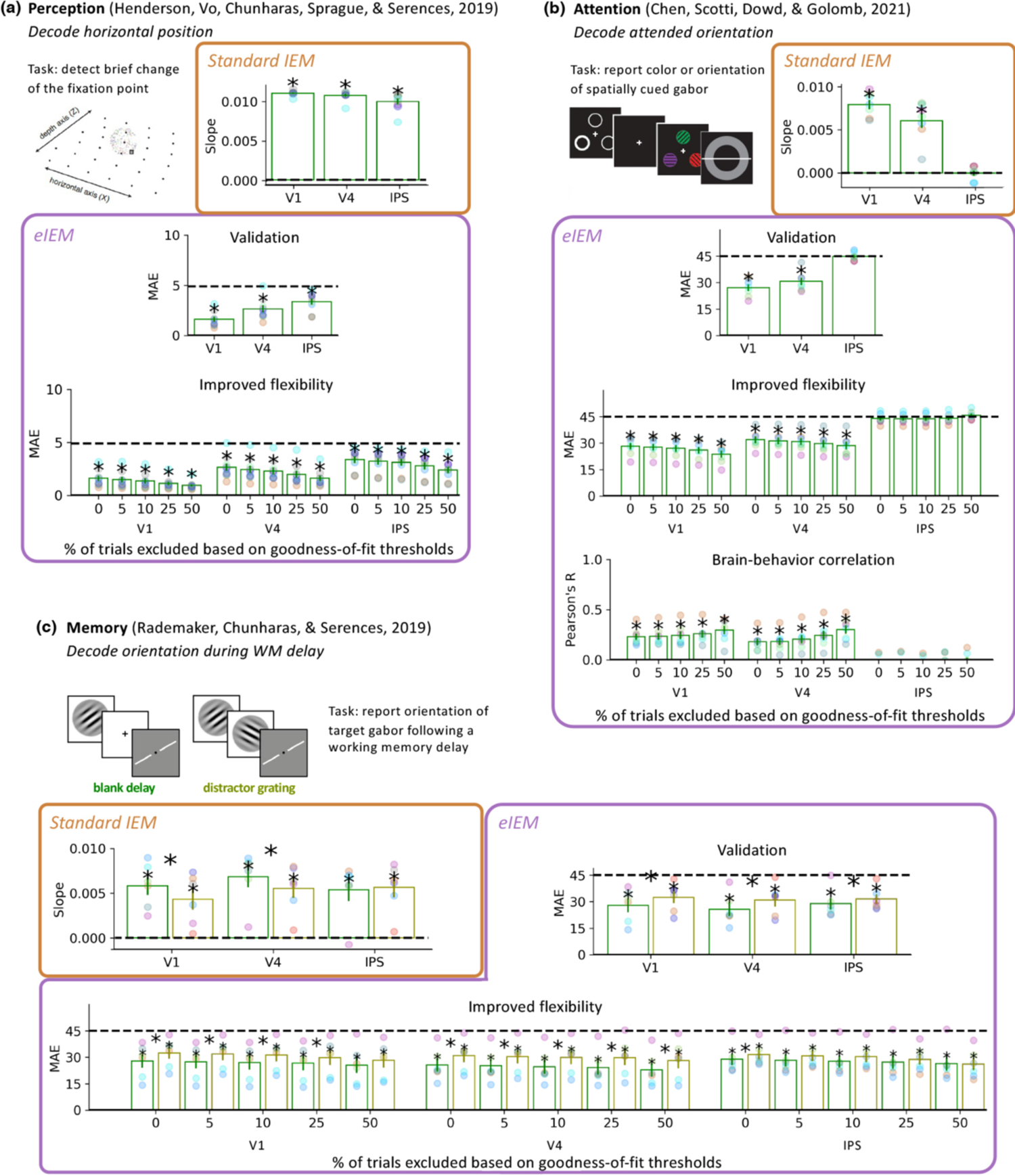
Application of eIEM to three real fMRI datasets spanning the topics of perception (a), attention (b), and memory (c). For each dataset, we show results obtained from a representative “standard” IEM align-and-average approach (orange boxes, see Methods for details), our eIEM procedure without thresholding (purple boxes, ‘Validation’ plots), and eIEM using increasingly stringent cutoffs based on goodness-of-fit (purple boxes, ‘Improved Flexibility’ plots). Bar plots depict the reconstruction quality, using average slope for the “standard” procedure (higher values are better) and MAE for eIEM (lower values are better) across subjects, with individual subjects overlaid as colored dots, for each of 3 ROIs (V1, V4, IPS). For the Attention dataset, the ‘Brain-behavior correlation’ plot additionally plots the trial-by-trial correlation between absolute behavioral error and absolute decoding error for each ROI and goodness-of-fit threshold. Error bars depict standard error of the mean, dotted black lines represent chance decoding, and asterisks represent statistically significant decoding (p<.05, no corrections for multiple comparisons). Overall results show that conclusions are similar between the “standard” align-and-average IEM approach and eIEM, but MAE is more interpretable (not based in arbitrary units) and not prone to the methodological concerns we discuss later in the Results. In addition, each dataset showed that MAE consistently improved with increasing exclusion thresholds, demonstrating the flexibility of goodness-of-fit to exclude noisy trials. Increasing exclusion thresholds also appeared to strengthen brain-behavior correlations in the Attention dataset. See Supplementary Table 1 for test statistics and p values.

In the Perception dataset^48^, we used both techniques to decode the horizontal position of a stimulus in V1, V4, and IPS. The “standard” align-and-average procedure revealed significant decoding performance in all 3 ROIs, with the strongest decoding (greatest slope) in V1, followed by V4, and then IPS. eIEM replicated this pattern. In the Attention dataset^47^, we used both techniques to decode the attended orientation within a multi-item, multi-feature stimulus array, in the same three ROIs. The “standard” IEM procedure revealed significant decoding in V1 and V4, but not IPS. eIEM again replicated this pattern. Finally, in the Memory dataset^43^, we used both techniques to decode the remembered orientation of a stimulus over two types of working memory delays: blank delay and distractor delay. With the “standard” procedure, the remembered orientation was successfully decoded in V1, V4, and IPS, with significantly greater decoding in V1 and V4 during the blank delay compared to the distractor delay. With eIEM, we replicated each of those results, with the additional finding of significantly greater decoding during the blank vs distractor delay in IPS.

### 2.2 Demonstrating the improved interpretability and versatility of eIEM

Having validated our method across three diverse fMRI datasets, we next use these same datasets to illustrate the practical advantages and additional features of eIEM.

#### Improved interpretability

eIEM lends itself to metrics that are easily interpretable and comparable across datasets due to decoding performance being measured in terms of prediction error. In contrast, the arbitrary units typical of the more standard aligned-and-averaged reconstructions are not easily interpretable. For example, in the Memory dataset, the “standard” procedure results in an average slope of .006 for the blank delay condition and .004 for the distractor delay condition in V1 (or cosine fidelity values of .100 and .098 as reported in the original paper^43^); eIEM replicates this pattern, but now with a more interpretable and meaningful metric: orientation can be decoded with an average error of 27.9 degrees in the blank delay and 32.5 degrees in the distractor condition. Note that the eIEM procedure can evaluate trial-by-trial reconstructions using any metric, not just the MAE metric, if a different metric better aligns with the researcher’s goals.

#### Incorporating the additional trial-by-trial uncertainty feature

Next, we tested the flexibility of eIEM to make use of the trial-by-trial goodness-of-fit information. The eIEM approach produces a best-fitting stimulus prediction and associated goodness-of-fit value (correlation coefficient) for each trial. It is important to emphasize that the correlation coefficient reflects the degree to which the reconstruction matches the *best-fitting* basis channel, not the basis channel centered on the correct stimulus. In other words, this goodness-of-fit information is obtained without circularity. It is obtained independently and prior to any calculation of prediction error.

To test the impact of using goodness-of-fit information on decoding performance, we performed an analysis where we excluded trials with the lowest 5%, 10%, 25%, and 50% of goodness-of-fit values (“Improved flexibility” subplots of Figure 2). This resulted in visible improvements in MAE (i.e., smaller decoding error) with increasing exclusion thresholds for all three datasets (linear regression revealed significant negative slope in all cases except for IPS in the Attention dataset). Notably, in the Attention dataset, MAE improved with increasing thresholds in V1 and V4 (where decoding was significant in the unthresholded analysis) but not in IPS (where decoding was at chance in the unthresholded analysis). Thus, the goodness-of-fit information can be used to improve decoding performance when a brain region contains reliable information about a stimulus, but does not produce false positives in the absence of observable stimulus-specific brain activity.

This trial-by-trial prediction uncertainty information could be used in several ways. One suggestion we put forth is that goodness-of-fit can be used to threshold reconstructions, such that worse-fitting trials may be excluded from analysis. This principle is analogous to the phase-encoded retinotopic mapping and population receptive field modeling techniques, where a set of models spanning the full stimulus space is evaluated for every voxel, and the parameters of the best-fitting model are selected as that voxel’s preferred stimulus, with the goodness-of-fit values then used to threshold the results^49–51^. Note that although r-squared is the more commonly used statistic for goodness-of-fit using regression, squaring the correlation coefficient loses potentially critical information about the sign of the correlation coefficient (e.g., a perfectly inverted reconstruction should *not* be assigned equal confidence as a perfect reconstruction in most cases), so we recommend the use of the r-statistic. For data scrutiny purposes or for experiments where an inverted reconstruction is theoretically informative, then researchers are free to adopt r-squared values or to not calculate goodness-of-fit values entirely. Alternative approaches to trial thresholding also exist, including weighting trials based on their goodness-of-fit (see Discussion).

Goodness-of-fits obtained using eIEM can also be applied more broadly, beyond the MAE decoding metric. That is, one could use eIEM goodness-of-fits to threshold trials while using any of the commonly employed metrics of slope, amplitude, cosine fidelity, etc. Goodness-of-fit thresholding can also be applied directly to (aligned-and-averaged) reconstruction curves. This could be particularly useful when one wants to use eIEM and quantify results with MAE, but also visualize reconstruction curves. We illustrate this point in Figure 3 using the above 3 fMRI datasets along with a simulated dataset. As increasingly strict goodness-of-fit thresholds were applied, the reconstruction curves were visibly improved. All of the quantification metrics similarly showed improvements with increasing goodness-of-fit thresholds. For the real fMRI datasets, these improvements generalized across all datasets, subjects (within-subjects and across-subjects), brain regions, and experimental conditions (mirroring the performance benefits shown in the “Increased flexibility” subplots of Figure 2). For the simulated dataset, these improvements generalized across the full range of simulation parameters tested, including number of trials, number of voxels, number and shape of basis channels, shape of voxel tuning functions, amount of noise, etc. A sampling of 78 eIEM simulations with randomly determined ground truth parameters across varying goodness-of-fit thresholds is depicted in Supplemental Figure 4, demonstrating the robustness of goodness-of-fit thresholding to improve reconstructions (code available to stimulate your own reconstructions on OSF: https://osf.io/et7m2/).

**Figure 3.**
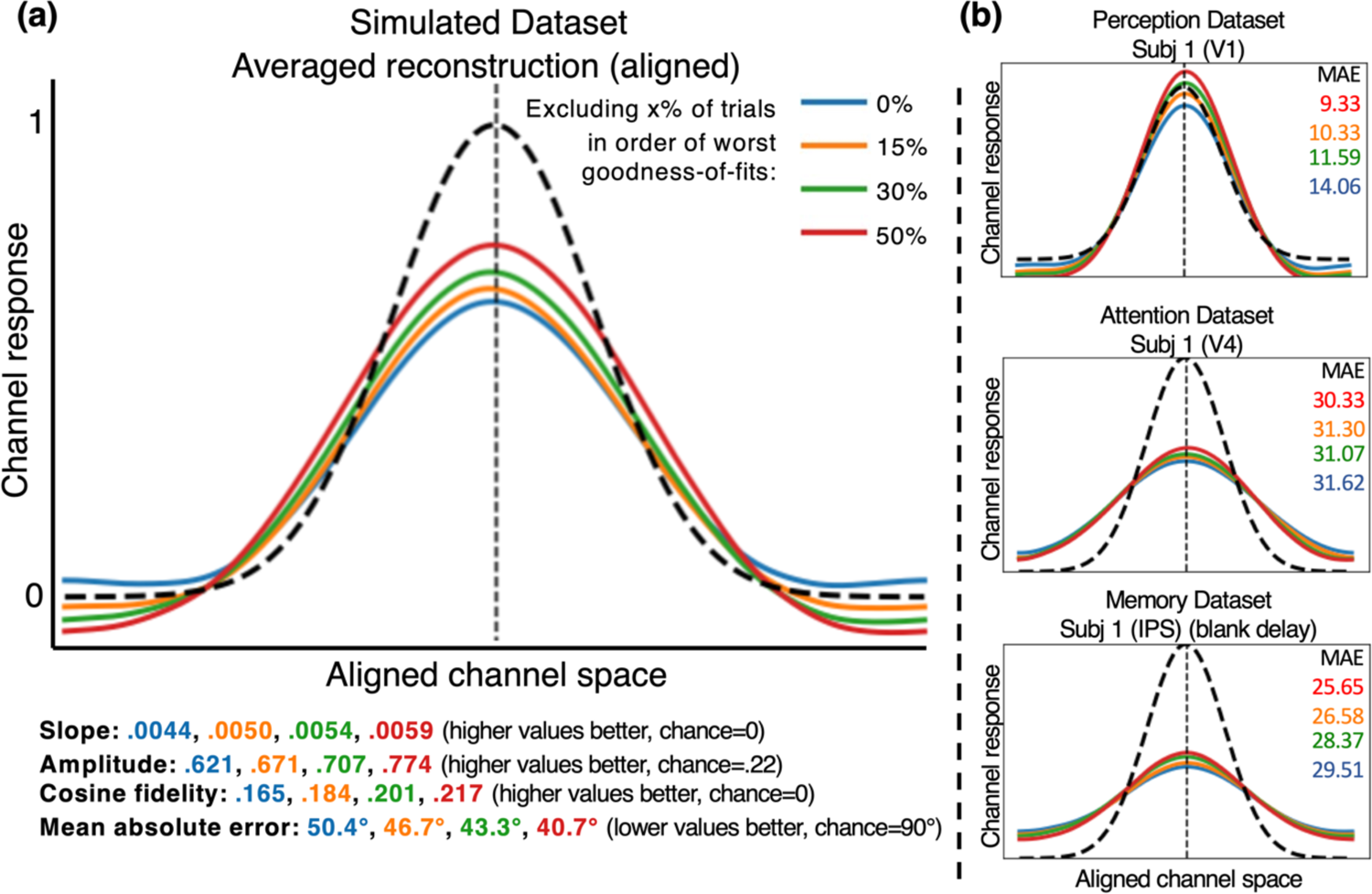
Simulated and real datasets show how eIEM goodness-of-fits can be applied to more traditional aspects of IEM, particularly visualization of reconstruction curves. Different colors reflect different goodness-of-fit thresholds, implemented by exclusion of the X% worst fitting trials. Dotted black reconstruction denotes the original basis channel at the aligned point (i.e., “perfect” possible reconstruction). Visual inspection of the reconstructions shows clear improvements with increasing thresholds. (a) Depiction shows the aligned-and-averaged reconstructions across 300 trials within a simulated brain region of 50 voxels, using a basis set of nine equally spaced channels. Several decoding metrics are shown below the curve, color-coded for each threshold level; all similarly benefitted from this trial thresholding, demonstrating the versatility of eIEM goodness-of-fits to benefit modeling outside of the eIEM framework. Results generalized across a wide variety of simulation parameters; a random sampling of 78 more eIEM simulations with randomly determined ground truth parameters are depicted in Supplemental Figure 4. Note that MAE typically, but not always, followed a monotonic function across thresholds. The code to reproduce these results and simulate reconstructions with varying parameters is available on OSF (https://osf.io/et7m2/). (b) We demonstrate the robustness of this approach to improve aligned-and-averaged reconstructions at the single-subject level across all three real fMRI datasets. Due to space limitations, we depict just three plots using the first subject of each dataset across three different ROIs, but this general pattern of stricter thresholds leading to improved reconstructions held across all datasets, subjects, brain regions, and experimental conditions.

#### Brain-behavior correlations

Finally, having trial-by-trial prediction error and goodness-of-fit values lends itself to analyses correlating neural measures with behavior. We demonstrate this in the Attention dataset (the Perception dataset did not collect behavioral responses, and in the Memory dataset behavioral performance was too close to ceiling [∼3° avg. error]). In V1 and V4, we observed a significant correlation between a trial’s behavioral error magnitude and neural prediction error. Moreover, the strength of these correlations increased with higher goodness-of-fit thresholds, as depicted in the “Brain-behavior” subplot of Figure 2 (Fisher z-transformed correlation test: V1: *t*(6)=6.85, p<.001; V4: *t*(6)=5.31, p<.001). We note that behavioral error itself did not noticeably change across these thresholds, suggesting that the goodness-of-fit information seemed to be reflecting noise at the level of the fMRI signal, not simply fluctuations in behavior or cognitive focus.

Altogether, these findings demonstrate several tangible benefits of using eIEM. Not only does eIEM produce robust, reliable results, but it can do so using a prediction error framework and metric that is more interpretable and intuitive. Moreover, the novel addition of trialwise goodness-of-fit values offers increased flexibility to improve both neural decoding power and brain-behavior correlations. Further, researchers can use the visualization approach depicted in Figure 3, where reconstructions are visually scrutinized alongside an overlaid perfect reconstruction, which we recommend to clarify interpretations and ensure data are scrutinized before relying on a numerical decoding metric.

### 2.3 Additional methodological concerns addressed by eIEM

The three real fMRI datasets analyzed above are useful validation cases because they contain robust findings (as may be more likely with published, publicly available datasets), allowing us to convey the improved interpretability and flexibility of eIEM in cases where the overall pattern of decoding is consistent with a “standard” align-and-average IEM approach. Crucially, however, there are also cases where we would expect eIEM results to diverge from the “standard” IEM results due to various methodological concerns inherent in the align-and-average approach. Below we illustrate some hypothetical cases susceptible to specific methodological concerns and limitations that are addressed by eIEM. These hypothetical cases are based on theoretical principles and illustrated by cherry-picked visualizations to showcase some ways that eIEM could meaningfully diverge from standard align-and-average steps; while it remains unclear how pragmatically relevant such divergences would be in real neuroimaging experiments, even less extreme versions of these patterns could produce misleading data via other IEM approaches.

#### Potential information loss due to the align-and-average step

Figure 4 depicts a few examples of how aligning and averaging reconstructions across trials could obscure important information. The hypothetical cases illustrate one case where every trial is accurately reconstructed (i.e., centered on the correct stimulus feature with minimal error, panel 4a) and one case where every trial produces inaccurate reconstructions (panel 4b). These two cases would be interpreted identically if a researcher were relying on standard aligned-and-averaged reconstruction metrics. In contrast, eIEM correctly identifies the first case as superior (lower MAE). Because eIEM evaluates prediction error on a trial-by-trial level and then averages the result, it avoids the pitfall of interpreting Figures 4a and 4b as reflecting the same quality of stimulus-specific brain signal.

**Figure 4.**
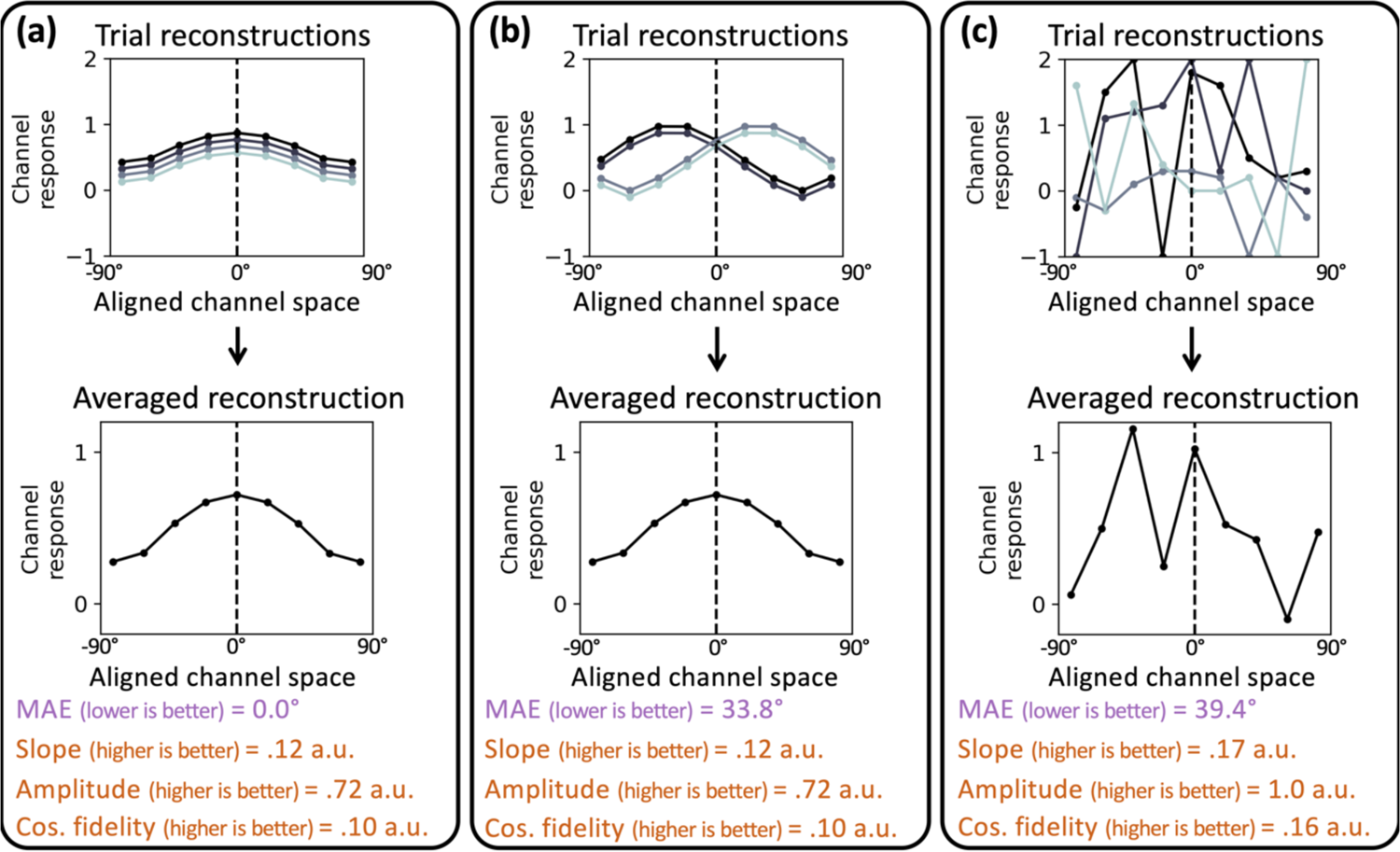
Visual depiction of how reconstructions with the standard align-and-average procedure can produce spurious results because of not considering the variability of trial reconstructions or the shape of the basis channel. For each of the 3 hypothetical examples (where data points were manually defined), the top row depicts four single-trial reconstructions, and the row below depicts the aligned-and-averaged reconstruction. In (a) each individual trial’s reconstruction accurately predicts the correct channel (i.e., the correct stimulus feature), appropriately reflected in the averaged reconstruction. In (b) each individual trial’s reconstruction predicts an incorrect channel. Averaging across trials leads to a misleading result, i.e., the standard align-and-average approach would consider (b) to reflect the same level of decoding performance as (a). In (c) each individual trial’s reconstruction is essentially noise, such that the averaged reconstruction results in a false peak around the aligned point; the “standard” procedure using align-and-average metrics results in spuriously superior decoding performance than both (a) and (b), with (c) having a higher amplitude, steeper slope, improved cosine fidelity, and narrower standard deviation when fit with a gaussian distribution. The eIEM procedure, calculating MAE from trial-wise prediction error, correctly concludes that case (a) shows the best decoding performance.

Another advantage of eIEM is that MAE is also less prone to bias from outlier trials compared to any of the align-and-average metrics. When using the align-and-average IEM approach, a single outlier reconstruction can disproportionately bias the shape of the averaged reconstruction, potentially completely flipping the averaged reconstruction in the most extreme cases. In contrast, the influence of an outlier is naturally capped for eIEM: Consider a hypothetical experiment composed of 300 trials where 299 trials predict the correct stimulus and one trial predicts the stimulus 180° away (assuming 360° stimulus space); because prediction error is calculated at the trial level and then averaged, an outlier trial could only increase MAE by a maximum 0.6°.

#### Falsely assuming a monotonic relationship between decoding performance and neural signal

In addition to providing preferred solutions to the potential concerns of outlier bias and information loss, there are other advantages to evaluating trialwise reconstructions using eIEM. As described earlier, our trialwise goodness-of-fit metric allows the researcher to predict which trials will likely produce better reconstructions (and hence better stimulus predictions). While other complementary approaches exist, for example using trial-by-trial decoding error to help extract top-down spatial signals^44^ or to correlate trial-by-trial decoding with memory-guided behavior across trials^24^, critically, eIEM’s approach to calculating trialwise predictions avoids another known issue.

As has been recently highlighted in the literature, IEMs produce reconstructions that depend on the choice of basis set^38, 39^. If the IEM procedure does not explicitly take this observation into account, this can result in faulty interpretations. Supplemental Figure 2 illustrates a rudimentary example of this issue that can be addressed with iterative shifting, but a more fundamental problem remains. That is, intuition – and standard practice – wrongly assume that a monotonic relationship exists between reconstruction quality measured by metrics such as slope, amplitude, and bandwidth, and the amount of stimulus-specific information in the brain signal. However, if the basis set consists of equally spaced and identical basis channels (which is typically the case), then a “perfect” reconstruction returns the exact shape of the basis channel^b^. Yet, the standard align-and-average IEM steps can produce reconstructions from noisy neural data that appear *better* than this “perfect” benchmark. We argue that it makes more sense to directly compare the shape of the reconstruction to the shape of the basis channel to make predictions and evaluations. The correlation table step we implement in eIEM automatically adjusts to consider the shape of the basis channel because it is the basis channel itself that is being used to obtain predictions, providing a more direct relationship between IEM performance and stimulus-specific brain signal.

Simply put, amplitude, slope, bandwidth, cosine fidelity, etc. are inferior metrics compared to the correlation table metric because they do not adapt to the choice of basis set. This is true even if a researcher were to skip the align-and-average step by evaluating reconstructions at the trial-by-trial level. For instance, using the amplitude metric, a higher amplitude at the aligned point is thought to reflect improved performance. If the basis channel ranges from 0 to 1, a perfect reconstruction should have an amplitude of exactly 1 at the aligned point, but reconstructions can feasibly have amplitudes far greater than 1. Likewise, the cosine fidelity (aka vector mean) approach of evaluating reconstructions by taking the dot product of the reconstruction and a cosine function could favor reconstructions with a wider width than expected under ideal conditions. Figure 4C illustrates an extreme example of this problem, where standard align-and-average IEM metrics would produce spuriously high values, potentially leading one to falsely interpret the neural signal from Figure 4C as producing a better reconstruction than the signal from Figure 4A.

Note that to avoid this pitfall, it is important to preserve the exact shape of the basis channel defined by the encoding model when fitting reconstructions. While some prior approaches partially account for the shape of the basis channel by, for example, fitting the aligned-and-averaged reconstruction with a curve of the same general shape as the basis function (e.g., Henderson et al., 2019^48^), such fitting procedures (1) operate on the aligned-and-averaged reconstruction, and (2) allow the model parameters to be flexible. This process could still produce misleading conclusions (e.g., fitting Figure 4c with a cosine and taking the amplitude would result in the appearance of superior decoding performance compared to Figure 4a). Meanwhile, eIEM evaluates the reconstructions at a trial-by-trial level, using a fixed basis channel shape (i.e., the only parameter that changes is the mean location in stimulus space). Our eIEM package implements this using the correlation table approach, which can be considered as a constrained version of curve-fitting, in the sense of optimizing the best location of the basis channel, but any approach that fits the trial-by-trial reconstructions via comparison to the exact basis channel shape used in the original encoder would also be appropriate.

## 3. Conclusions

Inverted encoding modeling has become a popular method for predicting stimuli and investigating neural representations because of its robust performance, simplicity of linear modeling, ability to predict untrained classes, and grounding in single-unit physiology. Our new eIEM technique operates under the same general scope of experimental scenarios as other IEM approaches while fixing important methodological concerns surrounding some standard IEM procedures, and offering increased functionality, flexibility, and interpretability of results. In other words, when it comes to whether significant stimulus-specific information can be reconstructed from a brain region, eIEM performs comparably to (i.e. at least as good as) other IEM approaches. The core advantages of eIEM are that it can also do more, increasing the types of analyses that can be conducted, as well enhancing the interpretability of results.

Namely, eIEM’s benefits are due to the following modifications: (1) iterative shifting of the basis channel returns reconstructions in stimulus space rather than impoverished channel space, (2) evaluating reconstruction quality using the same tuning function used to build the encoder is more theoretically sound than assuming a monotonic relationship between reconstruction shape and neural signal, (3) calculating trial-by-trial prediction errors can be used to evaluate in meaningful units when decoding stimulus features, improving interpretability (compared to an align-and-averaged reconstruction) and allowing comparison of decoding performance across different model specifications, and (4) calculating goodness-of-fits independently from stimulus predictions allows a trial-by-trial uncertainty measure that can be used to flexibly improve reconstruction quality and applications.

It is worth noting that while some prior papers have presented alternative IEM procedures that implement some of the modifications above, eIEM is unique in terms of its comprehensiveness and accessibility/simplicity. Moreover, the eIEM procedure is flexible: while our default eIEM procedure can be executed with a single line of code from our openly available software package, this code can be also modified to selectively adopt aspects of the eIEM pipeline while customizing others. For example, a researcher could choose to not to use the goodness-of-fit information, or to use measures other than MAE. Alternatively, a researcher could customize the basis sets or other model parameters to test more advanced questions.

We demonstrated the above advantages of eIEM using three fMRI datasets. Our validations demonstrated how our eIEM approach can be applied to both circular and non-circular stimulus spaces, is sensitive to variations in decoding performance across brain regions and experimental conditions, and can be used to accurately decode the contents of perception, attention, and working memory. Our modifications further allowed for decoding performance of each dataset to be directly compared to each other in intuitive units, allowed us to meaningfully link neural reconstructions with behavior, and demonstrated how uncertainty, measured via goodness-of-fit, can be leveraged to increase statistical power. Note that just because these three datasets produced consistent overall results (in terms of significance testing) across procedures does not ensure this will always be the case, as illustrated by the cases in Figure 4 where eIEM carries less of a risk of misinterpretation than other implementations.

One of the more novel aspects of eIEM is that researchers have the flexibility to exclude trials with noisier reconstructions based on our goodness-of-fit scores, which reflect how similar in shape each reconstruction is to the basis channel at the predicted stimulus. This flexibility can be used alongside *any* computational model or evaluation metric because researchers could first threshold trials using goodness-of-fits obtained from our eIEM approach before proceeding to implement non-eIEM steps. A core difference between goodness-of-fits obtained using eIEM compared to other IEM metrics that evaluate the shape of a reconstruction is that eIEM goodness-of-fits are obtained independently from prediction error—they are obtained without reference to the correct, ground truth stimulus (other metrics first center reconstructions at the correct stimulus and then evaluate the result, which is circular if used for trial thresholding). Note that we do not prescribe a specific cutoff for determining goodness-of-fit thresholds in this paper, rather, we simply offer that such an approach is possible for improving IEM performance. For example, a researcher could a priori decide to more heavily weight trials with higher confidences or simply exclude the noisiest 20% of trials. We depict examples of varying goodness-of-fit thresholds across a variety of simulation parameters in Supplemental Figure 3. We also stress that goodness-of-fit thresholds need to be determined before data collection or according to some sort of a priori, independent criteria.

Our method also improves interpretability by evaluating reconstructions in terms of prediction error. For example, “V1 showed 10° average prediction error and V4 showed 20° average prediction error” is more interpretable than “V1 showed .02 amplitude and V4 showed .01 amplitude” because the latter is in arbitrary units, whereas MAE is in meaningful units. Further, unlike amplitude or slope, the magnitude of prediction error can be directly compared to other experiments using the same stimulus space. This allows researchers to compare the quality of decoding across experiments or experimental conditions with differing model specifications.

In this paper we have referred to IEMs as a specific kind of encoding and decoding model that involves simple linear regression with population-level tuning functions. An advantage of eIEM is that improvements are accomplished without sacrificing this simplicity—the encoding model weights and the decoding model channel responses are simply estimated via least-squares estimation. There are more complex neuroimaging methods that can similarly be used to produce reconstructions via hypothesized tuning functions. For instance, Kay et al., (2008)^5^ decoded natural images from brain activity via voxel-level receptive field models that describe tuning functions across space, orientation, and spatial frequency. Naselaris et al. (2009)^3^ further produced Bayesian reconstructions of natural images via the combination of encoding models meant to estimate structural and semantic content. Van Bergen and colleagues^46, 52, 53^ introduced models where voxels with similar tuning account for shared noise and which produce trial-by-trial probability distributions such that uncertainty can be obtained similarly to our procedure (although the researchers discuss this in terms of testing Bayesian theories of neural computation rather than trial thresholding). Further work will be necessary to compare all these approaches, and we acknowledge that these more complex modeling approaches may be more useful than eIEM depending on the research question and the extent to which the researcher prefers model complexity over simplicity. For now, given the ample and broad interest in the cognitive neuroscience community towards implementing relatively more simplistic IEM procedures, we offer eIEM as an accessible means to improve reconstructions within the same familiar framework already popularized by IEM.

In summary, inverted encoding modeling has become increasingly popular in recent years, and yet the proper method for evaluating and interpreting IEMs has become increasingly uncertain. Our eIEM method is advantageous for both theoretical and practical reasons. From theoretical principles, we argue that eIEM is less susceptible to methodological and interpretational pitfalls than other typical implementations of IEM. We also demonstrate clear and practical advantages for evaluating reconstructions according to our eIEM method: as validated with real fMRI datasets, researchers can use eIEM to obtain improved reconstructions via thresholding with goodness-of-fits, compare decoding performance across experiments and varying basis sets using a metric grounded in meaningful units, evaluate performance in stimulus space rather than impoverished channel space, and obtain concrete stimulus predictions (with corresponding goodness-of-fits) for every trial rather than rely on a summary statistic based in arbitrary units. While there already exist approaches capable of addressing some of these concerns, our approach represents a suite of best practices that can be adopted by researchers in future work. Our procedure can be easily implemented with just one line of code using our accessible Python package (https://pypi.org/project/inverted-encoding; see Methods)^c^, and can be applied to investigate a vast range of research questions that involve reconstructing the contents of perception, attention, and memory from neuroimaging data.

## Materials & Methods

We performed analyses on both real and simulated data. For the real fMRI datasets, we used two publicly available published datasets^43, 48^ and one unpublished dataset from our lab^47^. Note that we only analyzed a subset of the data from each dataset, picking a single key analysis and analyzing one or two conditions across three brain regions for the sake of simplicity. The experimental paradigms and conditions / regions chosen are described more in each dataset’s respective section below.

### Inverted encoding model procedures

For all datasets, we performed a set of analyses using both a standard align-and-average IEM approach and the eIEM approach (as depicted in Figure 1, with the exception that we used iterative shifting for both approaches to facilitate more direct comparison). For both procedures, we used the same basis set. Basis channels can be modeled as

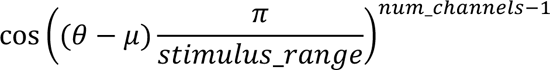

where θ is degrees in stimulus space, μ is the center of each channel, and *stimulus_range* is the range of stimulus space (e.g., 360° hues on a color wheel). The reasoning behind raising cosines to the *num_channels*-1 is to make the tuning curves narrower and more comparable to physiological findings^54^. Our eIEM Python package actually approximates this cosine with a gaussian function; this allows the user to experiment with different widths for their tuning functions via the optional parameter *channel_sd*. For the encoder, each voxel’s response was modeled as the weighted sum of the channels such that the observed trial-by-voxel fMRI activation matrix is equal to the dot product of the basis set and the weight matrix,

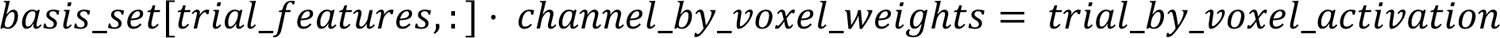

where *trial_features* is the feature (e.g., orientation) of the stimulus and *basis_set* is the matrix of channels with shape (*stimulus_range, num_channels*).

Once the weights matrix is estimated from the training dataset, it is inverted such that the weights matrix and the trial-by-voxel matrix are given and the channel responses (i.e., reconstructions) are estimated.

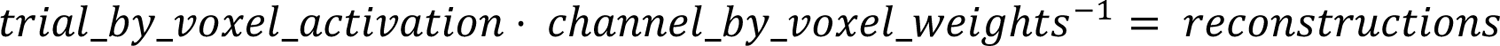

For all datasets, we used a basis set composed of nine equidistant channels each modeled as the gaussian approximation of 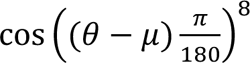. We used 10-fold (no shuffling) cross-validation, such that each iteration trained the model on 90% of the data and tested the model on the remaining 10%, repeated such that all trials were at one point decoded as part of the testing set.

For the ‘standard’ align-and-average approach, we aligned and averaged the single trial reconstructions into an average reconstruction and calculated slope as a traditional decoding metric. For the eIEM procedure, we calculated absolute prediction error for each trial via the correlation table metric and then calculated MAE. We performed these steps for each subject, ROI, and condition, and then we calculated the average slopes and MAEs across subjects.

For each condition and ROI, we assessed significance via permutation testing.

Significance tests were one-sided and uncorrected, calculated by comparing the t-statistic calculated from the actual data against the permuted null distribution of t-statistics (one t-statistic per each of 5,000 permutations). For eIEMs, we also repeated this analysis pipeline using varying levels of goodness-of-fit thresholds. That is, we discarded a certain percent of trials based on the worst goodness-of-fits and then calculated MAE using the remaining trials. The full list of t-statistics and corresponding p-values for the tests performed in Figure 2 are displayed in Supplementary Table 1.

### Perception dataset: Henderson, Vo, Chunharas, Sprague, and Serences (2019)

Data were obtained by downloading post-processed fMRI data associated with Henderson et al. (2019)^48^, publicly available on OSF (https://osf.io/j7tpf/). In this experiment, nine participants attended to a central fixation while a sphere (multicolored flickering dots positioned on the shell of a 3D sphere with radius 3.4°) was presented at varying positions along the horizontal and depth axes (depth achieved through stereoscopic MR-compatible goggles). The task was to detect a brief luminance change of the fixation point. Participants completed between 7 and 21 runs, where each run of 36 trials began with a sphere presented for 3s followed by a jittered intertrial interval (2-6s). There were also runs where participants covertly attended to the sphere, but we did not include these runs in the analysis. We only reconstructed horizontal position for simplicity and because position-in-depth was only sampled across six unique locations (varied sampling across the entire stimulus space is more appropriate for inverted encoding models) whereas horizontal position was sampled across 36 unique locations (from 0.9° to 9.8° eccentricity in both directions, collapsing across position-in-depth). We analyzed V1, V4, and IPS regions of interest which were defined via retinotopic mapping protocols where participants viewed rotating wedges and bowtie stimuli ^55^ while performing a covert attention task of detecting contrast dimming on a row of the checkerboard for the rotating wedge stimulus. We applied IEMs (following the procedures outlined earlier) to the post-processed data conducted by the authors of the original paper: Single-trial activation estimates consisted of averaged z-scored BOLD signal of the 3rd and 4th TRs following stimulus presentation. For more methods information, please refer to the original paper^48^.

### Attention dataset: Chen, Scotti, Dowd, & Golomb (2021)

Data were previously collected in our lab for another study Chen et al. (2021)^47^. In this experiment, seven participants completed a visual attention task. Each trial started with a central fixation cross. After 700ms, three circle outlines were displayed at equidistant locations surrounding the fixation cross for 200ms. One outline was thicker than the others, representing the spatial cue. Participants were instructed to covertly attend to the spatial cue location while maintaining fixation at the fixation cross. After 1100ms, three colored and oriented gratings were briefly displayed for 100ms, followed by a 200ms mask and a continuous orientation report. Participants were instructed to report the orientation of the grating that appeared at the location of the spatial cue.

There were also trials where participants were asked to shift attention to a different spatial location before the onset of the gratings, and entire runs where participants were asked to attend and report the color of the grating (instead of orientation), but we did not include these in our analysis. Participants completed at least 440 trials of each condition across multiple runs and sessions. We analyzed V1, V4, and IPS regions of interest: V1 and V4 were defined via retinotopic mapping protocols where participants viewed rotating wedges and bowtie stimuli^55^, while IPS was defined from the Destrieux atlas^56^ in Freesurfer (parcel labelled “S_intrapariet_and_P_trans”). To obtain single-trial neural activations for IEM, we modified a commonly used single-trial general linear model (GLM) approach^57^ to improve the model sensitivity and account for the large number of trials. Specifically, we conducted 40 GLMs per subject, where each GLM includes one regressor per run for one of the 40 trials in that run and one regressor per run for all the other remaining trials in that run. In this way, across the 40 GLMs, each trial in the experiment had an estimated single-trial beta weight. For more methods information, please refer to the original paper^47^.

### Memory dataset: Rademaker, Chunharas, and Serences (2019)

Data were obtained by downloading post-processed fMRI data publicly available on OSF (https://osf.io/dkx6y)^43^. We reanalyzed Experiment 1 where six participants underwent a visual working memory task. For each trial, a cue indicating the distractor condition was shown for 1.4s, followed by a target grating shown for .5s where participants were instructed to memorize its orientation, followed by a 1s blank delay, and then an 11s delay where 3 possible distractor conditions were possible: blank delay, Fourier-filtered noise, or distractor grating of a pseudo-random orientation.

Following an additional 1s blank delay, participants had 3s to report the orientation of the target grating, and finally a variable intertrial interval (3/5/8s). Each participant completed 108 trials per distractor condition. We reconstructed the remembered orientation based on the average activation patterns 5.6–13.6 s after target onset, as in Figure 1c in their paper, but only used the blank delay and distractor grating conditions for simplicity. To maintain consistency across procedures and datasets, we used the 10-fold cross-validation described earlier, with the blank delay and distractor grating conditions trained/modeled/evaluated separately from each other. We analyzed V1, V4, and IPS regions of interest which were defined via retinotopic mapping protocols where participants viewed rotating wedges and bowtie stimuli^55^. We applied IEMs to the post-processed data conducted by the authors of the original paper: Single-trial activation estimates consisted of averaged BOLD signal between 5.6-13.6s (7-17 TRs) after target onset. For more methods information, please refer to the original paper^43^.

### Simulated datasets

We used Python to simulate fMRI data from hypothetical brain region of interests with arbitrarily chosen numbers of voxels, where each voxel’s ground truth tuning function was the same shape as the basis channel. For Figure 1 and Figure 3, we injected random noise from a gaussian distribution into the trial by voxel matrix to produce noisier data as would be expected from a real fMRI experiment. Simulated fMRI data were subjected to the IEM procedures described above to depict the contents of Figure 1, Figure 3, and Supplemental Figures 2 and 4. Our Jupyter notebook on OSF (https://osf.io/et7m2/) contains Python code to reproduce Figures 1, 3, and Supplemental Figure 4, and can also be used to simulate reconstructions with varying parameters including number of trials, number of voxels, number and shape of basis channels, shape of voxel tuning functions, amount of noise, etc.

### Python package: inverted-encoding

We have released the Python 3 package “inverted-encoding” on PyPi (https://pypi.org/project/inverted-encoding/) and GitHub (https://github.com/paulscotti/inverted_encoding) for easy implementation of our eIEM procedure. The package contains two main functions, “IEM” and “permutation.”

For the “IEM” function, the only necessary inputs are an array of the stimulus features for every trial, the trial by voxel activations matrix (note: inputs other than voxels may be used for other modalities), the given stimulus range (e.g., stim_max=180 for stimulus values ranging from 0°-179°), and the number of folds to use for cross-validation. The basis set can be specified as an optional parameter and will otherwise default to a basis set composed of nine equidistant channels each modeled using gaussian functions that approximate 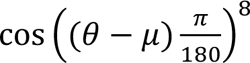. We use gaussian approximation because this allows the user additional flexibility to vary the width of the basis channel tuning functions via the standard deviation parameter. The stimulus space defaults to a circular space but can be optionally set to be non-circular via the Boolean parameter “is_circular.” We advise researchers to be careful regarding cross-validation, such that training and test data do not come from the same functional runs due to timepoints not being independent (autocorrelation, z-scoring, etc.). The final outputs are an array of each trial’s predicted stimulus, an array of each trial’s corresponding goodness-of-fit, each trial’s unaligned stimulus reconstruction, and each trial’s aligned stimulus reconstruction (aligned such that the correct stimulus is at zero in channel space). The user can then compute MAE (or any decoding metric that they prefer). MAE can be computed by averaging the (circular) absolute error between the predicted stimulus features and the actual stimulus features. We provide a convenience function, “circ_diff”, for computing circular error. The user can decide whether they want to threshold any trials using the provided goodness-of-fit values prior to calculating decoding performance.

For the “permutation” function, the only necessary input is an array of the actual stimulus features. For each iteration, the stimulus features are randomly shuffled and used as the predicted stimuli to compute the MAE. The function outputs a null distribution of MAE values for the user to compare against the MAE obtained from the “IEM” function. A more exact and computationally intensive method would be to rerun the entire IEM pipeline with shuffled stimulus labels on every iteration to obtain the null distribution. This can also be performed using our package by simply repeating the IEM function with a different shuffling of the stimulus features for every iteration. Our exploratory comparisons of null distributions obtained using both approaches across the three fMRI datasets discussed in the main text yielded no obvious differences.

## Data availability

The Perception dataset^48^ is publicly available on OSF (https://osf.io/j7tpf/). The Memory dataset^43^ is publicly available on OSF (https://osf.io/dkx6y). The Attention dataset^47^ will be made publicly available upon final publication of the original study; researchers may contact the authors to obtain this dataset before then.

## Code availability

Code to implement eIEM is available as a Python package (https://pypi.org/project/inverted-encoding). Code to reproduce Figures 1 and 3 are available on OSF (https://osf.io/et7m2/).

## Supplemental Information

### Supplemental figures

**Supplemental Figure 4.**
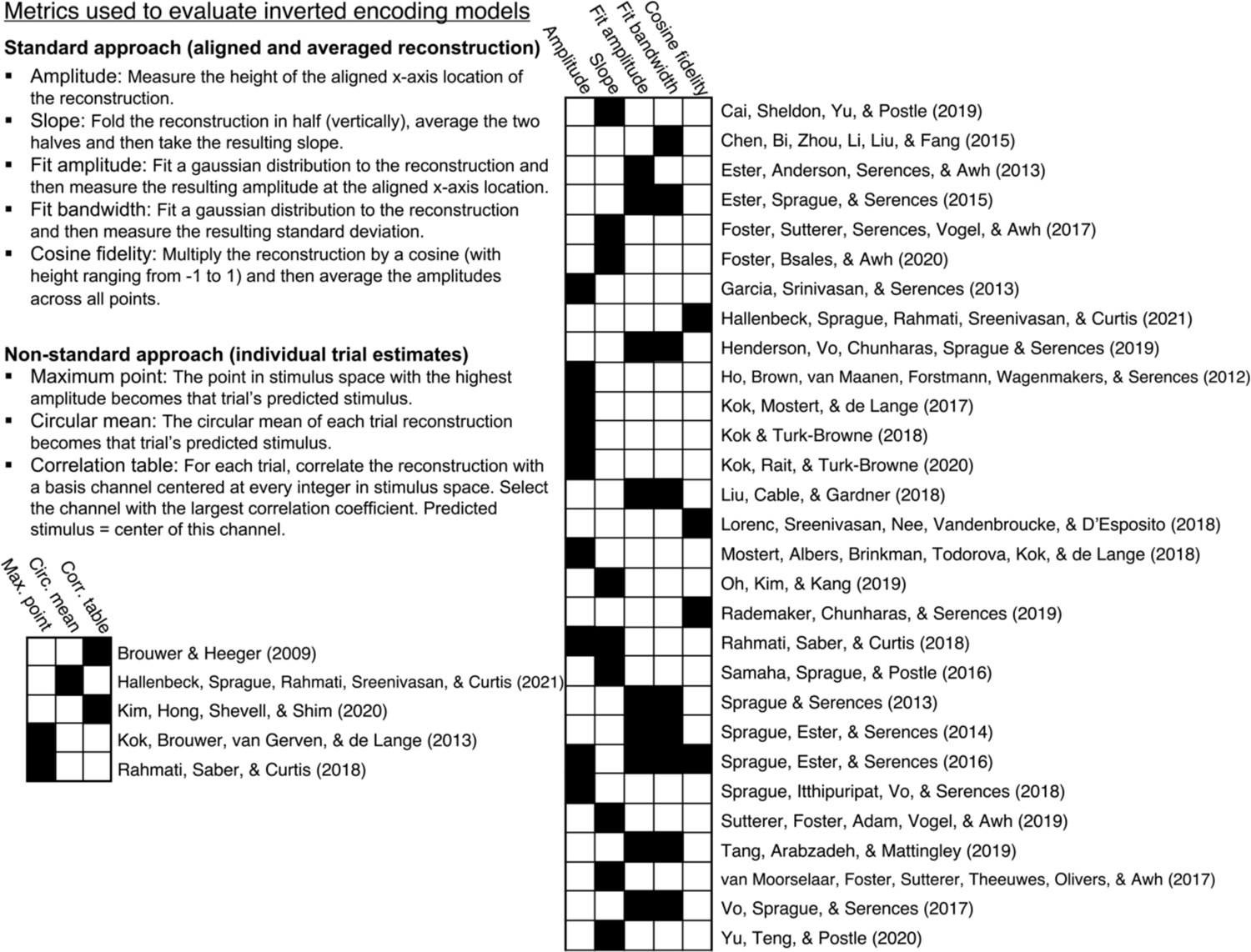
Summary of metrics used to evaluate IEM reconstructions in a sampling of published paper^1–31^. Note that the methodological concerns raised in the Results apply to the metrics labeled under the “standard” align-and-average approach. The non-standard approaches can be quantified with a single value (like the standard approach metrics) by taking the mean absolute error between predicted and actual stimuli. Note that not all papers used the exact same procedure to obtain their listed IEM metric(s), and that the reader should investigate the individual papers of interest for more exact methodological details.

**Supplemental Figure 5.**
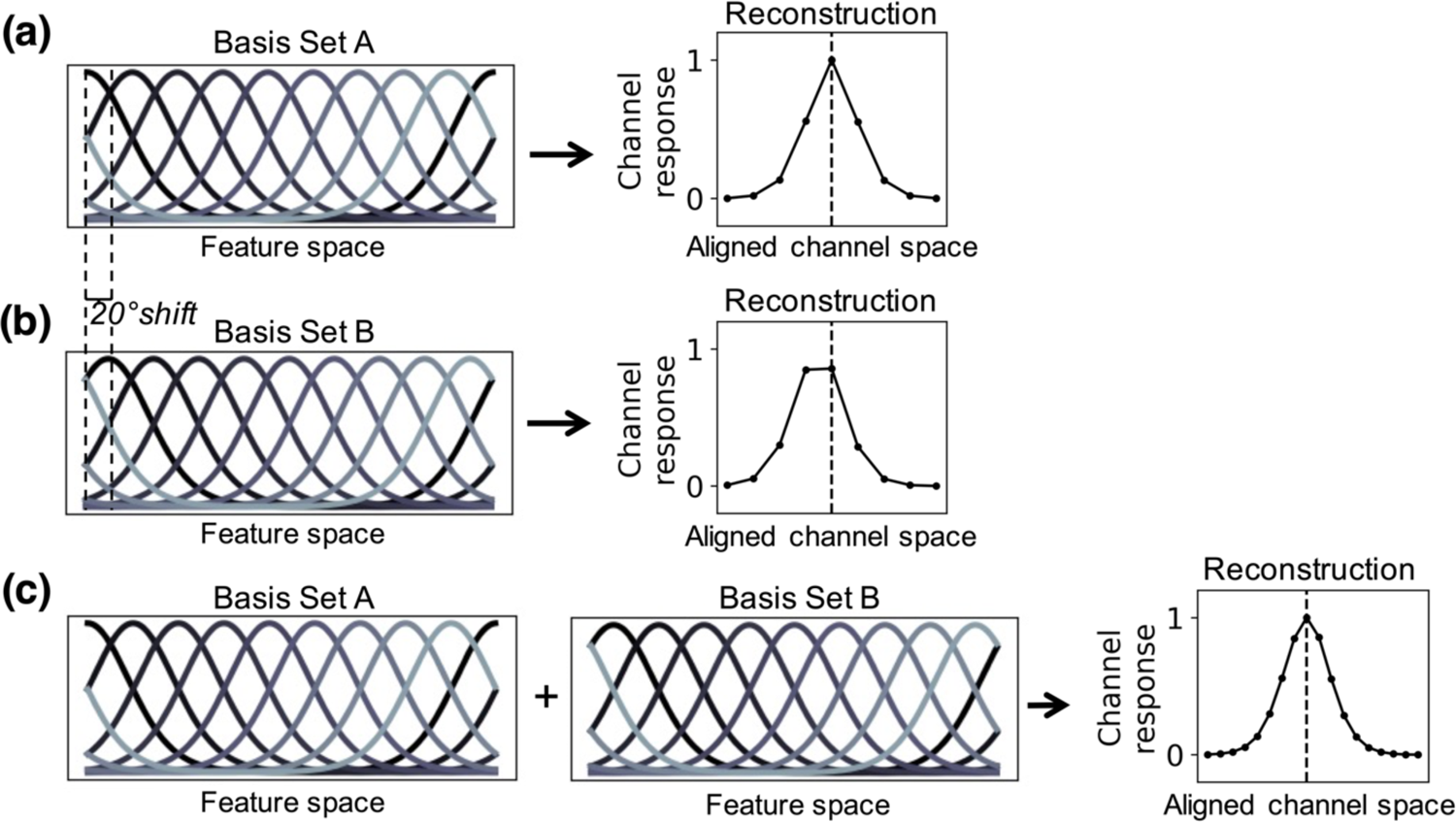
Simulation depicting iterative shifting and its benefits for IEM reconstructions. The simulated neuroimaging data (trial by voxel activations) are identical for all three cases shown, and the underlying voxel tuning functions for this hypothetical brain region are known (simulated ground truth). The trial by voxel activations were constructed to reflect perfect (zero noise) information with identical train and test sets, such that the resulting reconstructions should also be perfect. (a) Basis set happens to perfectly reflect the underlying voxel tuning functions (simulated ground truth). (b) Reconstruction of the same data, now with a slightly altered basis set (channel means circularly shifted by 20°). Note that in a real experiment, the ground truth actual voxel tuning functions are not known, so the experimenter’s arbitrary choice of channel centers could substantially alter resulting reconstructions, even if the same signal is present in all cases. This could result in misleading conclusions using the standard align-and-average IEM approach. (c) By combining the results of both basis sets, the channel space changes from *num_channels* to *num_channels*\*2, leading to a fuller reconstruction. This is the principle of iterative shifting: In eIEM, we repeatedly fit the encoding model with every possible (circular) shift of the basis set (i.e. 1° shifts) and then combine all of these iterations together. This produces a fuller reconstruction that is no longer impoverished by a limited number of *num_channels* points (i.e., the range of channel space becomes equal to the range of stimulus space) and allows the correlation table metric to be optimally applied. This iterative shifting procedure also aids more generally in producing more interpretable and less biased reconstructions, as it allows our decoding model to be capable of predicting any possible feature in stimulus space (that is, not solely the stimuli that are located at the centers of the basis channels). Note that iterative shifting does not change the fact that different basis sets result in different reconstructions, rather, it simply allows for the most accurate reconstruction given a set number of channels with defined bandwidths.

**Supplemental Figure 3.**
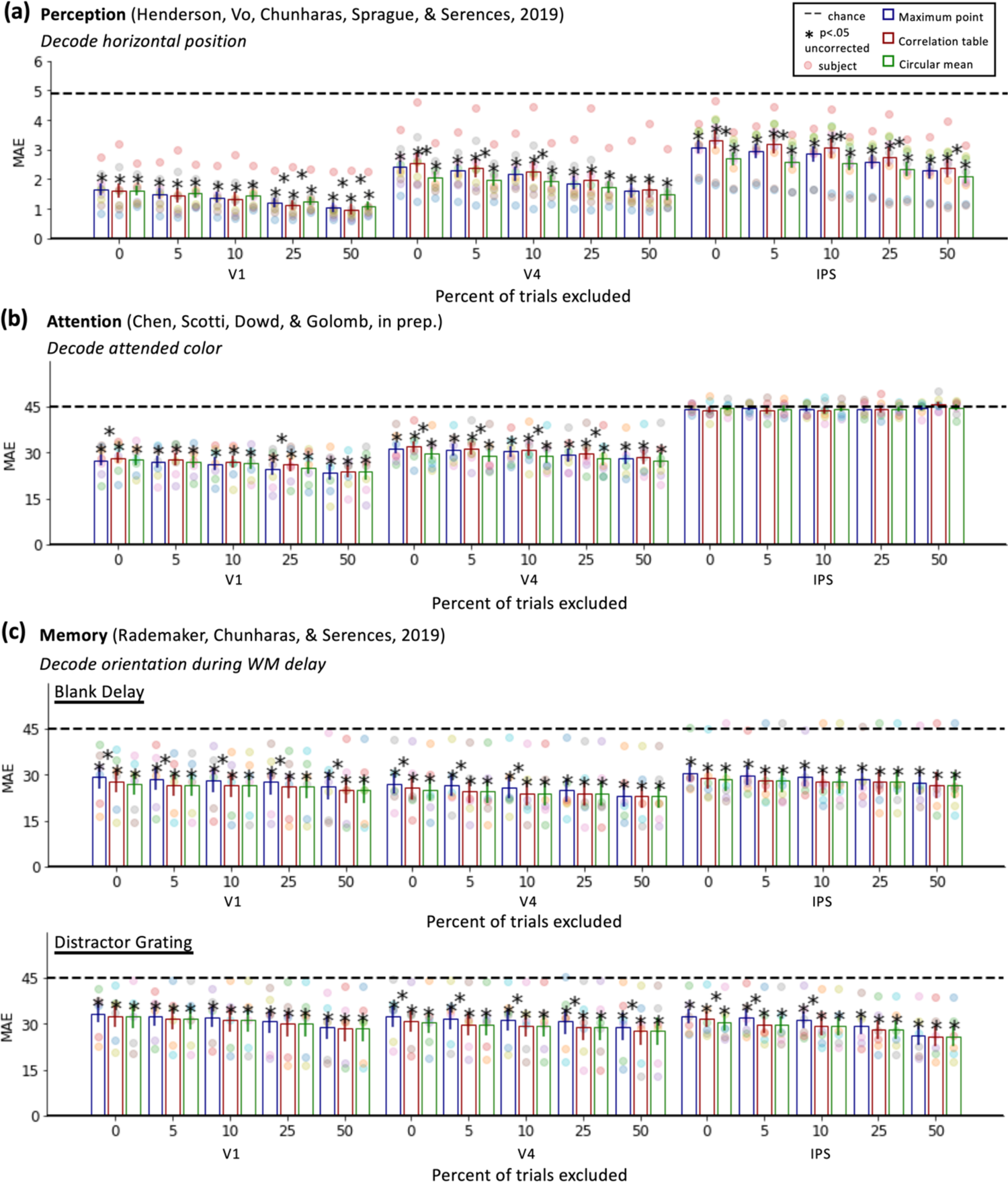
Comparison of MAE across varying goodness-of-fit thresholds using three possible techniques for calculating single-trial stimulus estimates (see Supplemental Figure 1): maximum point (blue), correlation table (red), and circular mean (green). In terms of improved MAE, “circular mean” appears to have performed best, but overall results are similar regardless of the single-trial estimation technique. As discussed in the main text, for the eIEM package we employ the correlation table measure based on the fact that it that accounts for the shape of the encoder’s basis channels and is parsimonious with goodness-of-fit calculations which rely on the correlation table approach.

**Supplemental Figure 4.**
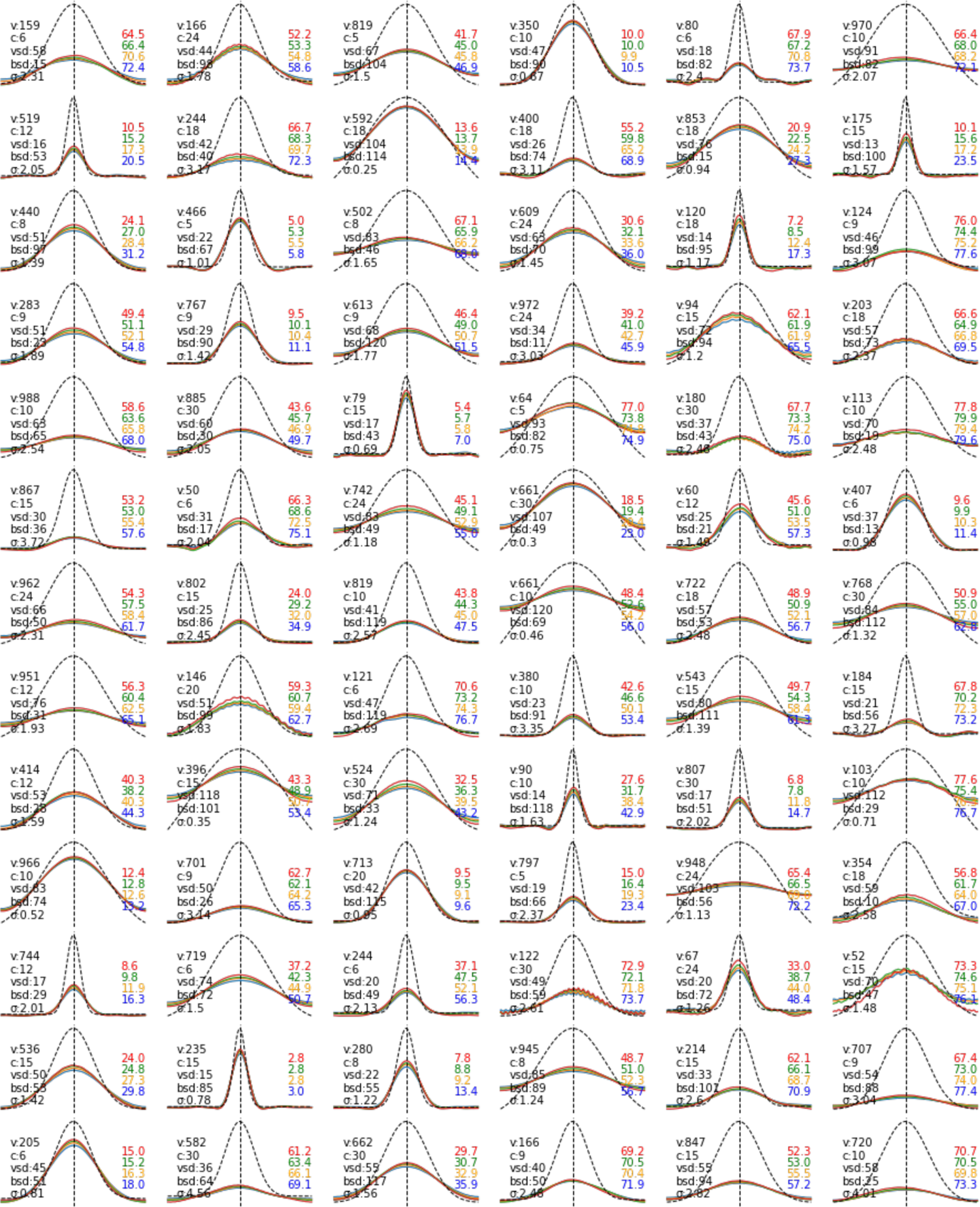
Random sampling of 78 eIEM simulations with varying ground truth parameters across 4 goodness-of-fit thresholds demonstrates the robustness of goodness-of-fit thresholding to improve reconstructions. Similar to Figure 3 in the main text, colored lines reflect different goodness-of-fit thresholds (red:50% green:30% orange:15% blue:0%), the dotted curved line depicts the basis channel (i.e., “perfect” reconstruction), and the vertical dotted line denotes the aligned point in channel space. Text on the left-side of each plot denotes the ground truth simulation parameters. “v” refers to the number of voxels, which was a random integer between 50 and 1000. “c” refers to the number of equally spaced basis channels, which was a random integer between 5 and 30. “vsd” and “bsd” refer to the standard deviation of the underlying voxel tuning functions (vsd) and basis channels (bsd), which were random integers between 10 and 120. “σ” was a random floating-point number between 0 and 5 and refers to the standard deviation of a normal distribution (mu=0) from which random noise was injected into the trial by voxel matrix. Colored text denotes the MAE for each thresholded reconstruction. Subplots were automatically removed and replaced if the reconstructions showed near-chance performance or near-perfect performance, where differences from thresholding would not be expected to be visible. Code to reproduce this figure is available on OSF (https://osf.io/et7m2/).

### Supplemental table

**Supplemental Table 1.**
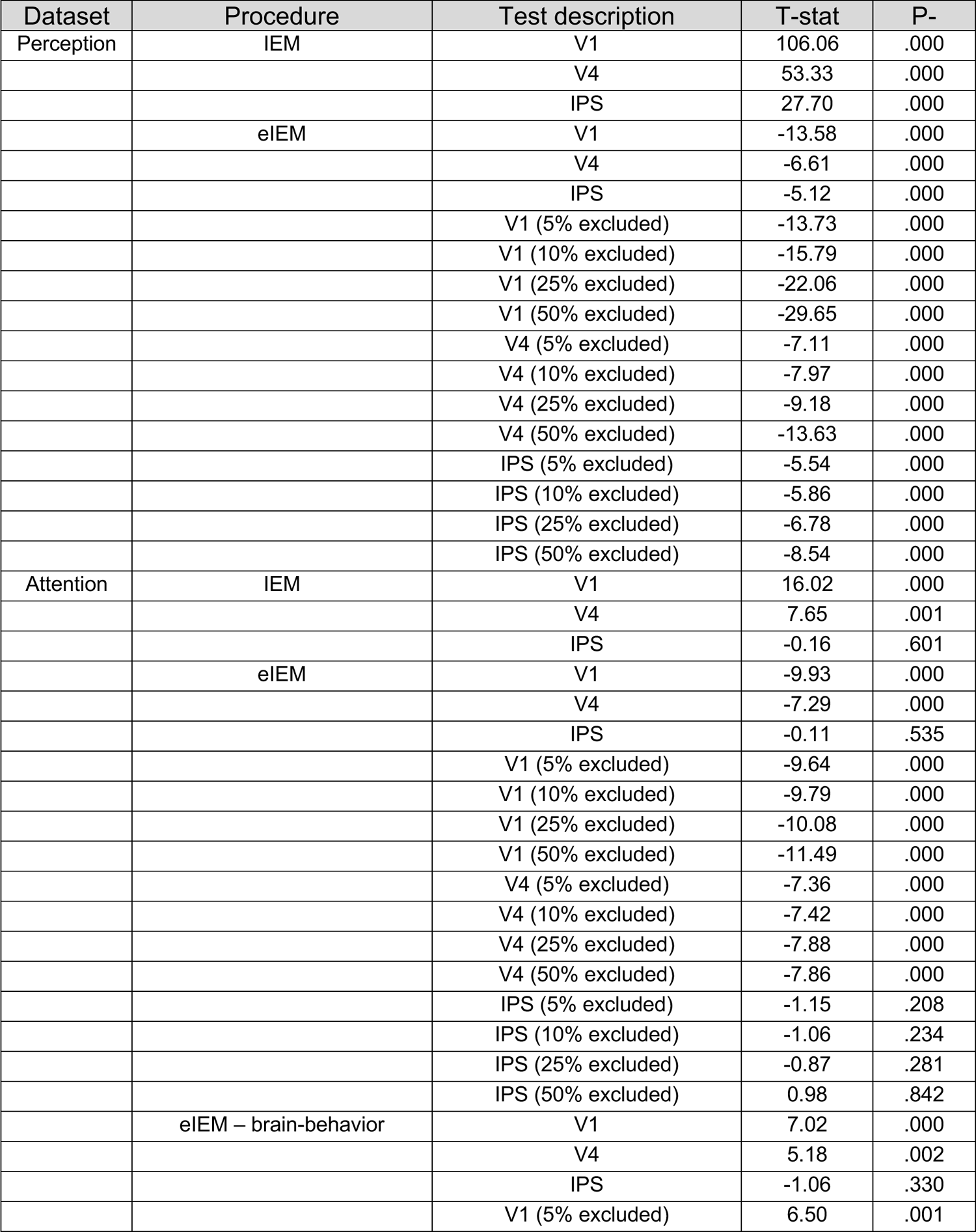
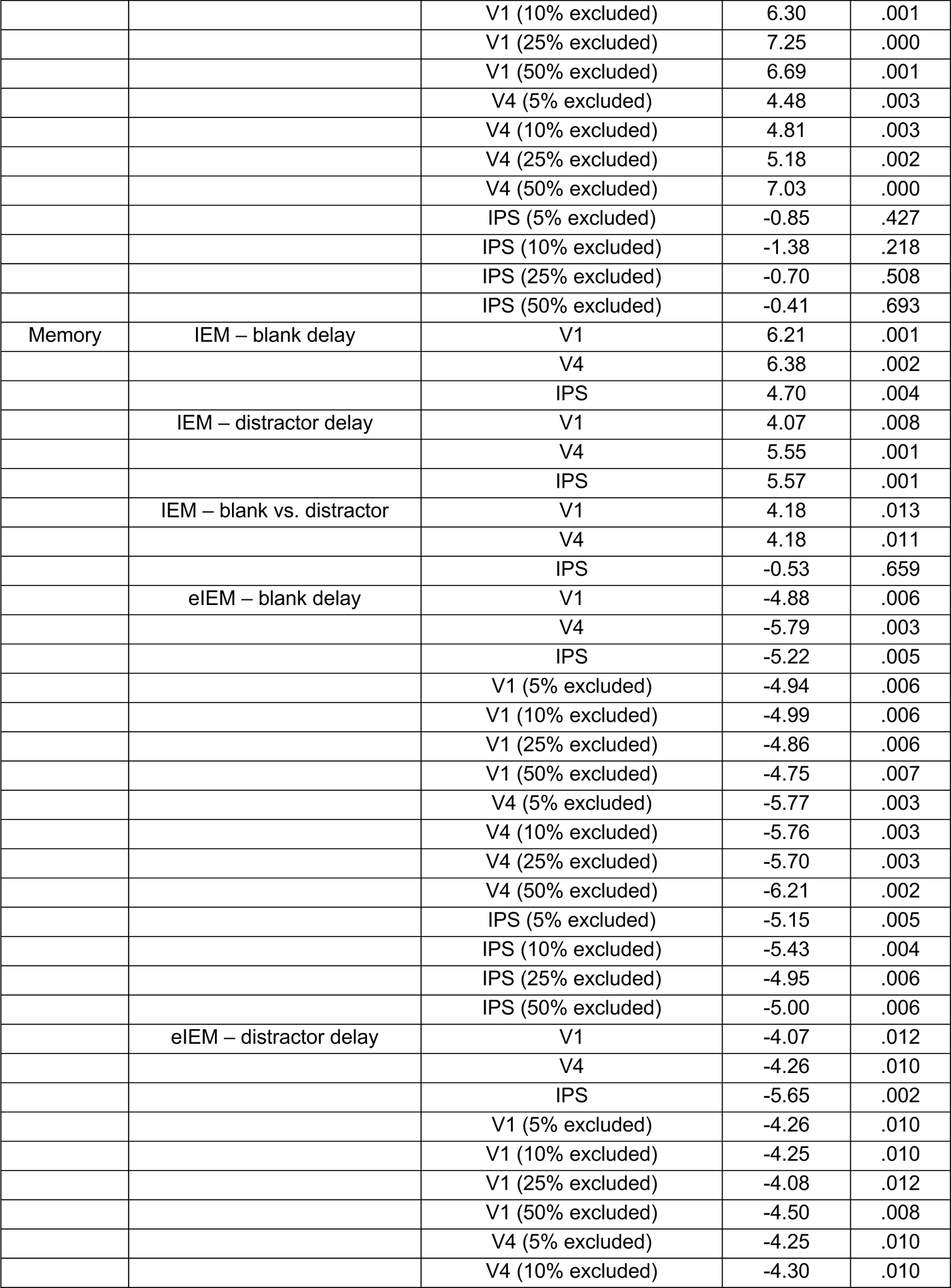
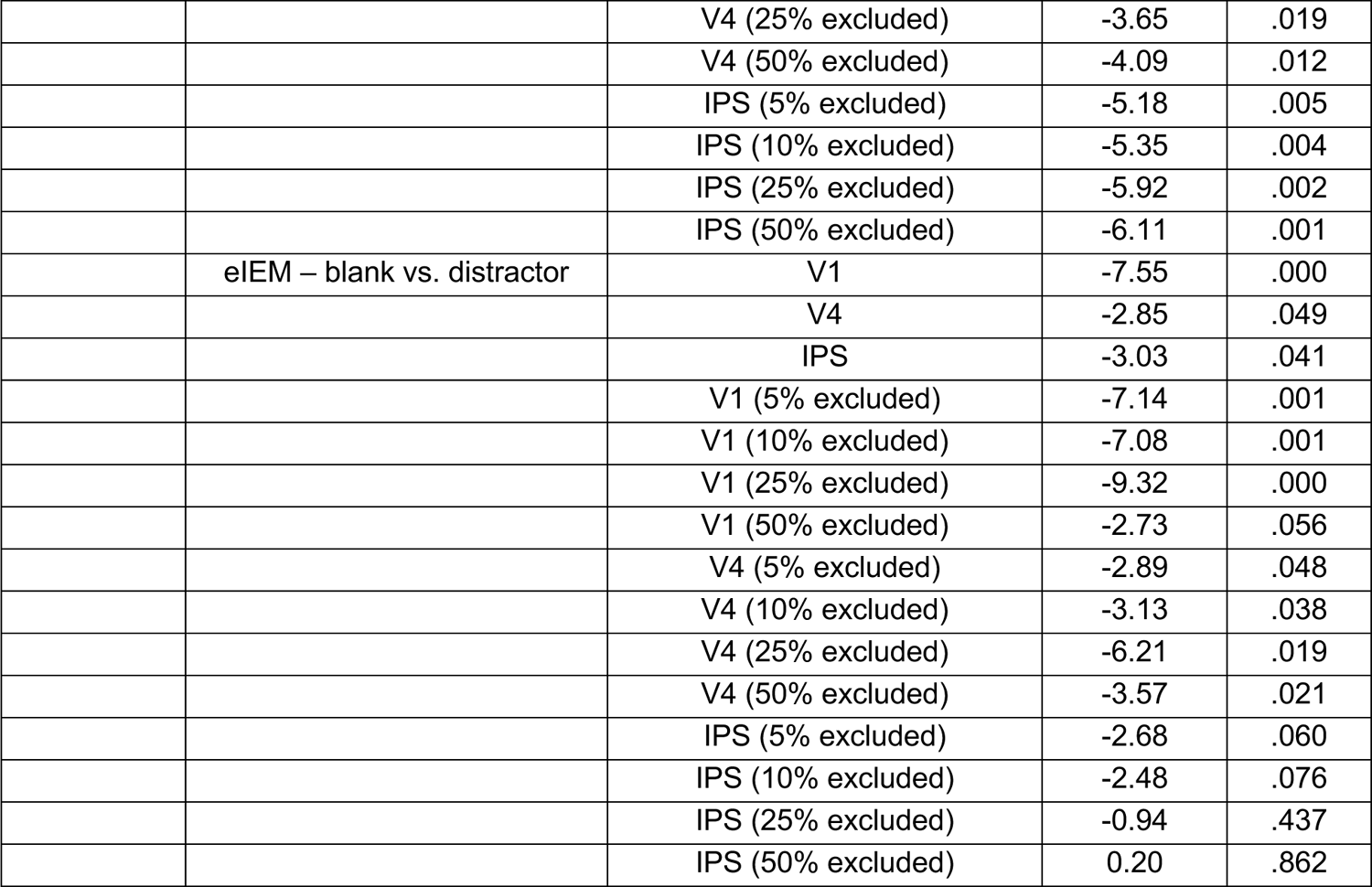
Statistics from tests depicted in Figure 2 from the main text.

## Footnotes

a There are other techniques besides the correlation table approach that have been used to obtain trialby-trial stimulus predictions within the IEM framework, which we refer to as “non-standard approaches” in Supplemental Figure 1. We directly compare mean absolute error calculated via these different nonstandard approaches using real fMRI datasets in Supplemental Figure 3. No approach was conclusively better than the others, and we picked the correlation table based on the fact that it that accounts for the shape of the encoder’s basis channels and is parsimonious with goodness-of-fit calculations which rely on the correlation table approach, as described below.

b There are some cases where the basis channel is not representative of the perfect reconstruction. For instance, if training a model with intermixed experimental conditions, or if using basis channels that are not identical or are unequally spaced. In these cases, one can simulate what a perfect reconstruction looks like at each stimulus feature and use these tuning functions in the same manner as the correlation table approach. We also note that there may be cases where visual comparison of reconstructions is the goal and numerical reconstruction evaluations are unnecessary.

c Note that while our Python package greatly simplifies inverted encoding modeling, we caution users to still scrutinize and visualize their data (e.g., averaged reconstructions can sometimes falsely look good as depicted in Figure 4, and reconstructions should always be visualized before relying on numerical decoding metrics).

